# Inhibitory Evoked Potentials as a Spatially Dependent Intraoperative Marker of Clinical Tremor Reduction

**DOI:** 10.64898/2026.08.24.746616

**Authors:** Zoe Paraskevopoulos, David Crompton, Sarah Iskin, Houyou Fan, Suneil K. Kalia, Mojgan Hodaie, Andres M. Lozano, Luka Milosevic, William D. Hutchison, Jurgen Germann, Milad Lankarany

**Author notes:** Correspondence to: Milad Lankarany Krembil Brain Institute, University Health Network, 60 Leonard Ave, 4KD512, Toronto, ON M5T 0S8, Canada.

## Abstract

Deep brain stimulation (DBS) of the ventral intermediate nucleus (Vim) of the thalamus may be used to treat medication refractory essential tremor. Using recordings from in vivo human Vim neurons, our previous work has suggested that evoked potentials (that we termed quasi-evoked inhibition) ∼2 ms following high frequency microstimulation pulses may be related to inhibitory synapses onto the Vim. Here, we investigate whether (***i***) quasi-evoked inhibition is related to clinical tremor reduction, and (***ii***) if quasi-evoked inhibition is dependent on the stimulation location within the Vim. By developing an objective determination of the presence or absence of quasi-evoked inhibition and utilizing accelerometer recordings, we showed that recordings with quasi-evoked inhibition at 100 Hz microstimulation exhibit greater tremor reduction than those without (P < 0.05, BF > 30). The number of stimulation pulses with quasi-evoked inhibition is also correlated with tremor reduction (rho = 0.18, P < 0.05) at all stimulation frequencies >=100 Hz. Furthermore, by analyzing microelectrode trajectories reconstructed from structural MRIs, we found that proximity to the ventral caudal border (P < 0.005) and to a previously established sweet spot (P < 0.05) are anti-correlated with the number of stimulation pulses with quasi-evoked inhibition. Our findings suggest that quasi-evoked inhibition is a potential biomarker of tremor reduction by means of network inhibition, and the more posterior regions of the Vim may allow for better recruitment of inhibition. This may be useful for closed-loop stimulation design.

## Introduction

Essential tremor (ET), is the most common tremor disorder, affecting 0.32% of the global population.^1^ It is thought to be a disorder of the cerebellum and previous research has demonstrated reduction of inhibitory receptors in the dentate nucleus of the cerebellum compared to healthy controls.^2,3^ This lack of inhibition may be propagated to the cortex, which is mediated by the motor thalamus. Since the Ventral intermediate nucleus (Vim), or the motor thalamus, is crucial in relaying activity from the cerebellum, it is often used as a target for invasive deep brain stimulation (DBS).^4^ High frequency (> 100Hz) DBS aims to suppress Vim’s neuronal firing and reduce tremor severity.^5^ However, the exact mechanism of Vim-DBS is not fully understood. Some evidence suggests that the recurrent activation of targeted synapses leads to synaptic depression.^6^ Others have proposed network inhibitory recruitment as a DBS mechanism, or a combination of both hypotheses.^7^

Using computational techniques and experimental Vim-DBS data, we proposed that without increased network inhibition, the model could not be fit to clinical Vim microelectrode data.^8^ Additionally, we found that while all recorded neurons exhibited evidence of synaptic depression at high frequencies, in a subgroup of neurons an evoked potential that emerges after ∼50 pulses that may be related to network inhibition.^7^ These neurons with “quasi-evoked inhibition” (QEI) are significantly more suppressed overall and temporally when the evoked potential is present. Since the Vim is an excitatory thalamic nucleus that projects mainly to primary motor cortex and thalamic interneurons, we previously hypothesized that the excitatory input induced from DBS allows for the recruitment of inhibition.^7,9,10^ Thalamic interneurons reciprocally inhibit the Vim, and the thalamic reticular nucleus receives glutamatergic inputs from the cortex but then has GABAergic projections back onto the Vim.^9,11^ However, the relationship between this scenario and clinical outcome remained unexplored. In this work, we hypothesize that QEI is a biomarker of improved clinical outcomes in DBS therapy for tremor reduction.

Furthermore, while there is research demonstrating “sweet spots” to target within the Vim for optimal tremor reduction, imaging and stereotactic targeting does not lead to implantation of contacts exactly at these targets reliably.^12^ Since QEI is only observed in a subpopulation of Vim neurons, we hypothesize that it is spatially dependent and corresponds with previous research identifying Vim “sweet spots”.^12,13^ To test our hypotheses, we evaluated QEI against accelerometer tremor reduction and utilized MRI imaging to determine what regions could be attributed to high observance of evoked potentials.

## Materials and methods

### Human experimental data

12 people with essential tremor or dystonia undergoing either Vim-DBS implantation surgery or Vim surgical thalamotomy participated in this study. Demographic and clinical data are included in Supplementary table 1, and surgical methods are described in supplementary methods. A triaxial accelerometer was used to record the scalar sum of accelerations during tremor. Recordings were taken from the wrist contralateral to stimulation. Baseline recordings were taken until a sustained tremor could be recorded, and then the recording continued for the full extent of the stimulation train. MRI imaging was available for 7 people (Supplementary table 1).

### Detection and classification of QEI

Once artifacts were removed, QEI events could be localized. The event classification criteria includes (Figure 1):

**Figure 1.**
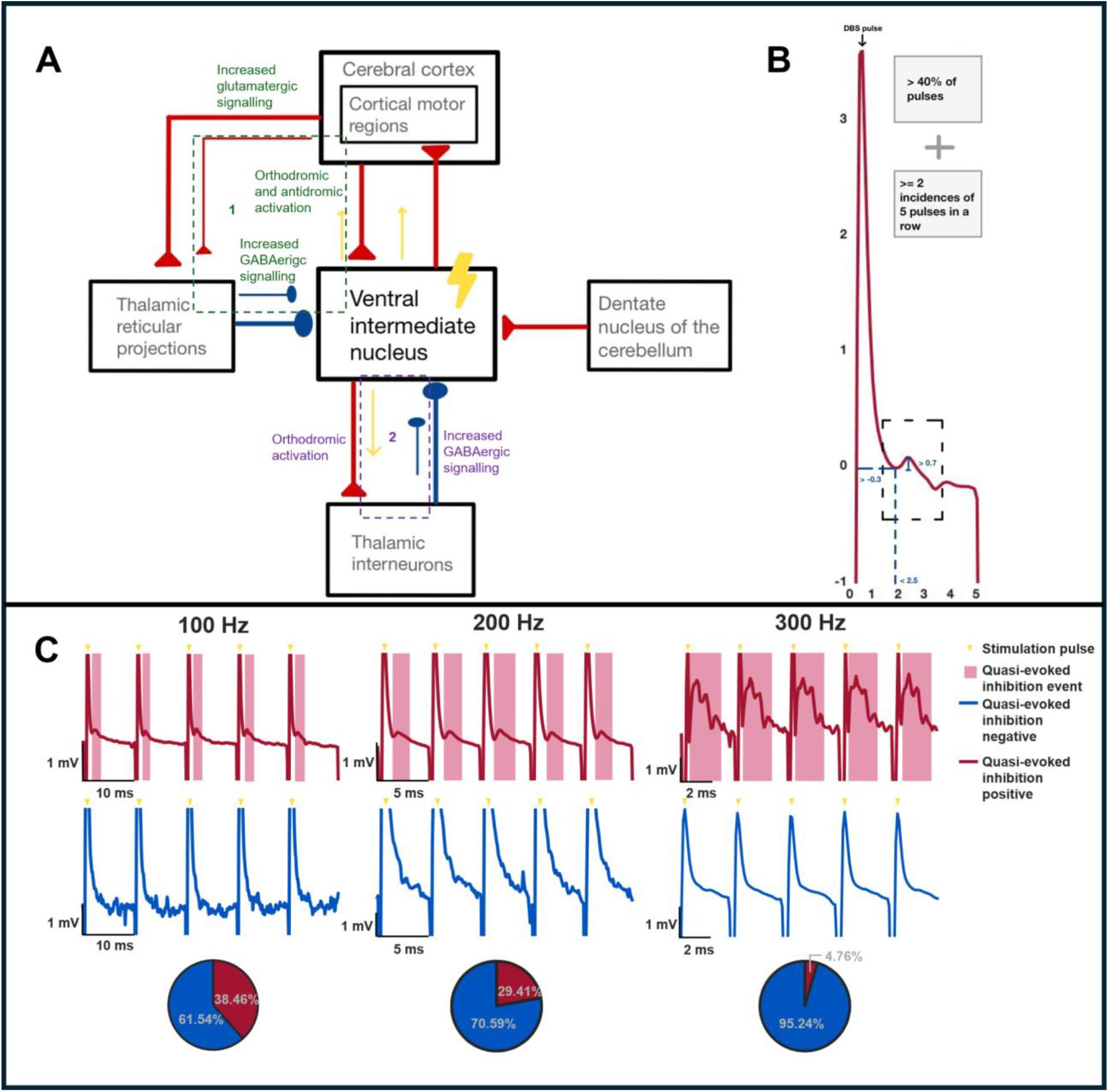
Proposed mechanism of QEI and classification requirements: Subfigure **A** shows the direct afferent signals that project onto the Vim as well as efferent outputs. Excitatory signals are depicted in red and inhibitory signals in blue. The two proposed mechanisms of QEI are numbered. The first proposed mechanism (surrounded by green dotted line) involves stimulation orthodromically and antidromically activating the cortex. This leads to increased glutamatergic signaling to thalami’s reticular projections which ultimately leads to increased GABAergic signaling to the Vim. The second proposed mechanism (surrounded by purple dotted line) involves orthodromic activation of inhibitory thalamic inter neurons, which cyclically inhibit the Vim in response. Subfigure **B** shows the classification criteria of the positive-going evoked potential, QEI. It must begin within the first 2.5 ms, have an amplitude greater than 0.7 mV, and not immediately follow a spike. Additionally, for a whole recording to be considered QEI positive, it must follow at least 40% of DBS pulses and have 2 or more incidences of 5 consecutive pulses in a row with QEI. Subfigure **C** shows examples of quasi evoked inhibition and the absence of QEI in samples at all three frequencies. The pie plots demonstrate the portion of the total recordings deemed QEI positive at each frequency.

1. The start point of the QEI event occurs before 2.5 ms into the inter-pulse interval.
2. The QEI event had an amplitude greater than 0.7 mV.
3. The starting voltage of the QEI event was greater than -0.3 mV.

Once all QEI events were localized, stimulation trains were classified as “QEI positive” if they met the following criteria (Figure 1):

1. There were at least two occurrences of five consecutive stimulation pulses with QEI following.
2. QEI events followed at least 40% of stimulation pulses.

More information about detection of QEI can be found in supplementary methods.

### Tremor analysis

For tremor reduction from baseline, the accelerometer recordings during the stimulation train, and the same time duration of stimulation immediately prior to the first pulse were analyzed. The accelerometer recordings were bandpassed between 3-10 Hz using a sixth order finite impulse response filter. For the baseline segment, the root mean squared amplitude of the signal was computed. Since there is a delay in tremor reduction after the onset of stimulation, for the stimulation segment, root mean squared (RMS) amplitude was computed in 0.2 sec windows. The window with the minimum RMS amplitude was identified, and the RMS amplitude was computed from that time point until the end of the segment to account for the return of tremor before stimulation ceased which was observed in some recordings. To compute the final tremor reduction from baseline, the ratio between RMS amplitude during stimulation and RMS amplitude during baseline was taken and subtracted from one. QEI spectral plots were computed from 100-250 Hz in steps of 1 Hz, using window lengths of 100 ms and steps of 10 ms. QEI spectral plots were visualized using the FieldTrip toolkit.^14^

### Structural MRI evaluation

To analyze the relationship between the distance on the trajectory to the ventral caudal (Vc) border and percent QEI, the first location where the patient exhibited paresthesia was noted and designated as the Vc. It was then determined the distance, in mm, between the Vc border and any other points marked for microstimulation during the implantation.

T1w MRIs were analyzed using Lead-DBS software.^15^ Statistical parametric mapping was used to linearly coregister pre-operative and post-operative MRIs.^16^ Pre- and post-operative scans were then normalized to MNI space using advanced normalization tools.^17^ Coregistration and normalization were manually reviewed and repeated using FSL software^19^ linear image registration tool^18^ and nonlinear image registration tool^17^ if deemed poor. Brainshift was corrected in postoperative acquisitions by applying a refined affine transform calculated from pre- and post-operative subcortical areas. DBS electrode trajectories were reconstructed manually by both Z.P and H.F. The coordinates of the Vim boundary were extracted as defined by Ewert et al. (2017)^20^ and the most caudal point where it intersected with each trajectory was defined as the Vc border in MNI space. The rest of the MNI coordinates were manually added. To assess the distribution of evoked activity of the Vim, barycentric interpolation was performed and mapped onto a grid overlaying the Vim. Gaussian kernel density estimation was done to assess the probability of a sampling at each point and used as a mask to remove data from under sampled regions. Visualizations were generated using Plotly.^21^

### Statistics

Comparisons were made between QEI positive and negative groups 100 and 200 Hz frequencies for tremor reduction. One-tailed Wilcoxon rank sum tests were conducted with Bonferroni corrections of alpha for two comparisons. Furthermore, Bayes factor computations were computed from the Mann-Whitney U test, using the <u>Distribution-Free Bayesian Analysis</u> package in R.^22^ To assess the correlation between percent QEI and percent tremor reduction, a Spearman correlation and a P-value computed from a t-test was used since data was monotonic. Additionally, to plot the correlation line, a linear regression was completed.

The effects of depth of trajectory and Euclidean distance to the location of the sweet spot defined by Papavassilou et al. (2004)^12^ and mapped to MNI space^15,17^ by Horn et al., (2017) ^15^ was assessed on QEI percentage and clinical tremor reduction. A random patient intercept linear mixed effects model was fitted with fixed stimulation localizations. Significance was assessed by fitting the null model, permuting the residuals, and adding the fitted null values to re-fit the model. This was done 5000 times, and a two-sided p value was computed based on the number of slopes fitted under the null hypothesis were greater than the observed slope.

## Results

### QEI is associated with an increased tremor reduction from baseline

We observed the highest prevalence of QEI during 100 Hz stimulation (38.46%, n_positive_ = 10, n_negative_ = 26), then 200 Hz stimulation (29.41%, n_positive_ = 20, n_negative_ = 68), and only one sample at 300 Hz stimulation (4.76%, n_positive_ = 1, n_negative_ = 21). We compared tremor reduction from baseline between QEI and positive and negative recordings for 100 Hz stimulation and 200 Hz stimulation (300 Hz stimulation excluded as only 1 recording displayed QEI). After Bonferroni adjusting p-values for multiple comparisons, we found strong evidence of increased tremor reduction in QEI positive recordings at 100 Hz stimulation (*Z* = 177, *P* = 0.026, BF_10_ = 65.12) and moderate but insignificant evidence of increased tremor reduction in QEI positive recordings at 200 Hz stimulation (*Z* = 935, *P* = 0.23, BF_10_ = 6.39) (Figure 2B). Furthermore, we found that percent QEI and percent tremor reduction were weakly correlated (rho = 0.18, 0.15 - 0.71, *P* = 0.038) (Figure 2C).

**Figure 2.**
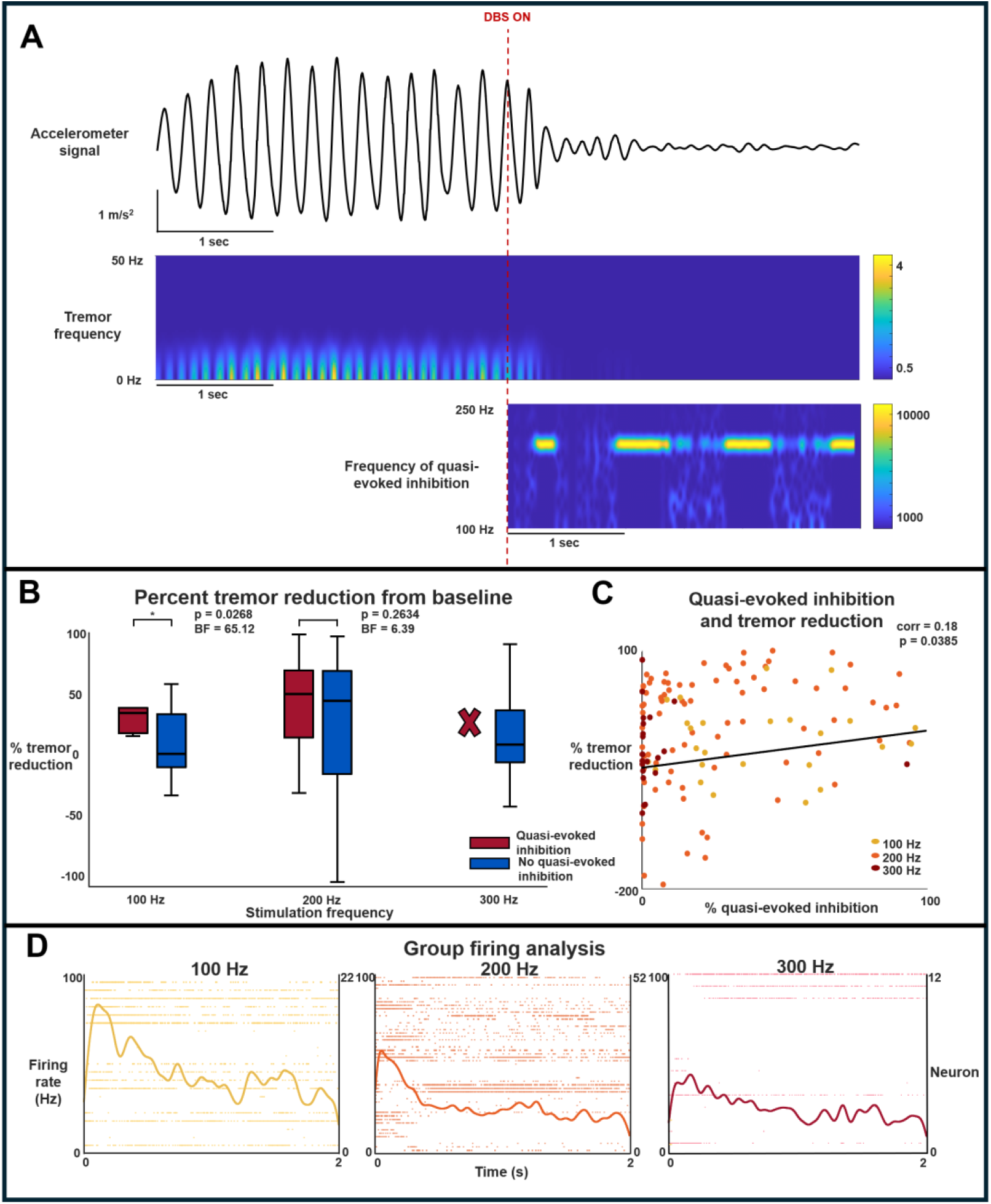
Increased tremor reduction in response to high-frequency stimulation associated with QEI: Subfigure **A** shows an example of a raw tri-axial accelerometer signal experiencing tremor at baseline and its silencing in response to 200 Hz microstimulation. Additionally, a spectrogram depicting the time-frequency power of QEI during the same stimulation period as the accelerometer recording. Subfigure **B** depicts the percent tremor reduction compared to baseline following microstimulation at 100 Hz (left), 200 Hz (middle), and 300 Hz (right). Each plot is divided into samples with QEI (red) and without QEI (blue). P-values and bayes factor values comparing tremor reduction with and without QEI are listed. No plot for tremor reduction with QEI at 300 Hz is shown because the sample size is only one. Subfigure **C** plots the percent of DBS pulses with QEI following for each sample against the percent tremor reduction from baseline following stimulation. Samples with 100 Hz stimulation are plotted in yellow, 200 Hz plotted in orange, and 300 Hz plotted in red. The line determined from a linear regression is plotted in black and the Spearman correlation and associated p-value are listed. Subfigure **D** shows the raster plots and associated firing rate plots using a 5 ms kernel for all neurons at each frequency. Mann-Whitney U test: *\* p-value < 0*.*025*.

**Figure 3.**
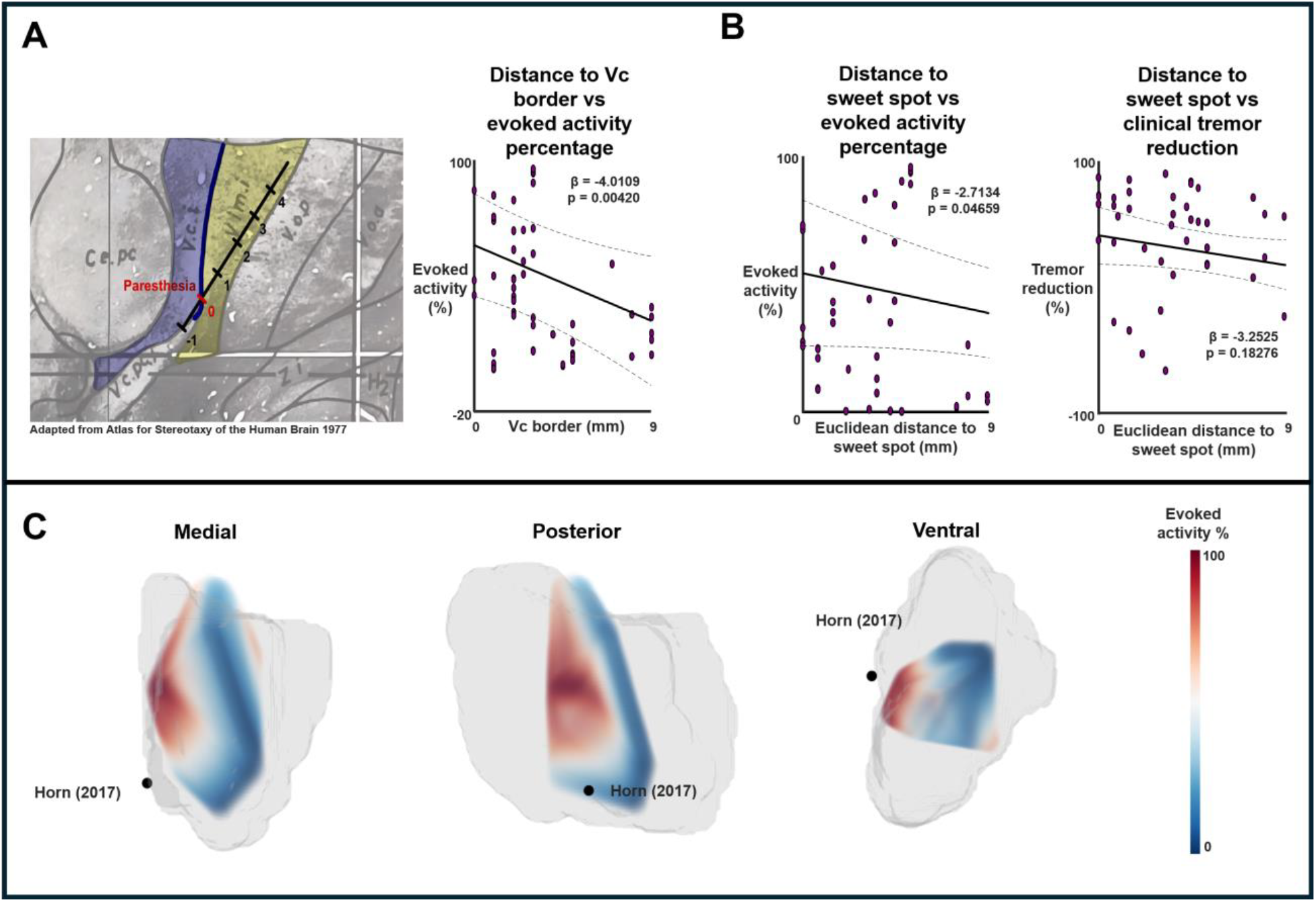
QEI is grouped ventrally and caudally: Subfigure **A** is adapted from the Schaltenbrand– Wahren brain atlas: Atlas for Stereotaxy of the Human Brain (1977) to show the anatomical border between the Vim (yellow) and Vc (purple) and how samples are taken and measured along the trajectory. A linear mixed effects model is used to show the relationship between distance from the Vc border and evoked activity percentage. Subfigure **B** shows the underlying Vim rendering (grey) along with the distribution of evoked activity in MNI space as a heatmap. This is shown from medial, caudal, and ventral views and the sweet spot as defined by Horn (2017) is plotted in black. Subfigure **C** shows linear mixed effect models between Euclidean distance to the sweet spot as defined by Horn (2017) and evoked activity percentage, and a linear mixed effect models between Euclidean distance to the sweet spot as defined by Horn (2017) and clinical tremor reduction. All linear mixed effect model beta and p-values are displayed.

### QEI appears strongly at the ventral and caudal borders

QEI percentage significantly decreased as the distance from the VC border increased (β = ™4.0109, 95% CI ™5.681 to ™2.3408; permutation p = 0.00420). Additionally, close distances to the sweet spot defined by Horn et al., (2017)^15^ was significantly associated with high percentages of QEI (β = ™2.2088, 95% CI ™3.7835 to -0.6341; permutation p = 0.04659). Interestingly, while close distances to the sweet spot in MNI spaces was associated with increased clinical tremor reduction, it was not significant (β = ™3.2525, 95% CI ™5.5888 to -0.9162; permutation p = 0.18276).

## Discussion

In this work, we developed an objective methodology to detect QEI events and determine whether a Vim single unit extracellular potential recording is “QEI positive” in general. Previous research has correlated field evoked potential amplitudes in other DBS nuclei to successful clinical outcomes.^23^ This work is the first evidence that QEI is a neurophysiological biomarker of clinical tremor reduction.

In comparison to the literature, our overall tremor reduction values (Figure. 2) were slightly lower but close to the microstimulation results in Milosevic et al 2018.^5^ However, when looking at the QEI group stimulated at 100 Hz, the overall tremor reduction, 33%, was much higher than the 17% observed in Milosevic et al. 2018.^5^ We hypothesize two explanations for this observation. Paraskevopoulos et al. 2026 found a larger difference in cellular inhibition between groups with QEI and without QEI at 100 Hz than 200 Hz.^7^ This could explain the larger clinical tremor reduction difference during the 100 Hz condition, as cellular inhibition is also positively correlated with tremor reduction.^5^ Another possibility is that inhibitory structures are being recruited but not firing at a high enough rate to induce evoked potentials at over 40% of stimulation pulses at 200 Hz.

Furthermore, we found that QEI is frequency dependent even within high frequencies. We found only one sample out of over 20 at 300 Hz that was QEI positive and the remaining samples had less than 5% of stimulation pulses associated with a QEI event. This was not a clinically effective microstimulation frequency compared to the QEI positive group at 100 Hz (33% reduction) or both 200 Hz groups (49% and 43% reduction) as the tremor reduction was only 7%. Previous research has shown that evoked potential oscillations in the subthalamic nucleus are dependent on clinically effective stimulation frequencies.^24^ This might explain why 300 Hz stimulation had a low tremor reduction, as it is not engaging the inhibitory network. It also leaves the questions whether clinically optimal stimulation frequencies (130 and 180 Hz) may maximize inhibition recruitment; or if stimulation frequencies resulting in worsening tremor (5 or 10 Hz) lead to even less Vim inhibition than baseline. Since the excitatory burst is not reliably produced in the 300 Hz microstimulation, this further supports our previous hypothesis that it is necessary to recruit inhibition.^7^ Birdno et al. (2011) found that constant DBS results in distruption of cerebellar tremor bursts, but DBS at the same frequency with intermittent pulses did not reduce tremor.^25^ The positive correlation between tremor reduction and number of pulses with QEI may suggest that even after inhibition is recruited, it must be maintained to disrupt cerebellar tremor bursts.

Next, we found that a higher percentage of QEI was associated with closer proximity to the Vc border. This aligns with previous research as Milosevic et al. (2018) found that closer proximity to the Vc border was predictive of greater tremor reduction.^5^ Since this work showed a correlation between QEI and clinical tremor reduction, it is not surprising that this effect would also carry over to the spatial relationship with the Vc border. Additionally, when the distribution of evoked potentials was mapped over the Vim, the highest concentration of stimulations with high percentages of evoked activity were located on the posterior surface of the Vim. Previous research has demonstrated in primates that the ventralis posterior lateralis pars oralis, a more posterior structure associated with the motor thalamus, is the main termination site of cerebellar afferents and provides the most projections to the hand controlling area of M1.^26^ Since pathological oscillations implicated in ET are projected from the cerebellum^27^, stimulating a location with a high concentration of these projections should be ideal. Previous research has demonstrated that targeting the dentato-rubro-thalamic tract also reduces tremors.^28^ Given that this also not a part of the basal ganglia circuit, future works may investigate whether QEI can also be observed at this target.

## Limitations and future directions

The QEI determination was based on percent of stimulation pulses with an associated evoked potential, but this was not adjusted for stimulation frequency. As mentioned previously, inhibition might be engaged, but not at a frequency high enough to result in QEI at greater than 40% of pulses at 200 Hz stimulation frequency. Many 200 Hz samples were close to the 40% threshold used to classify QEI positive recordings. In future works, the QEI algorithm may be tuned to account for differences between stimulation frequencies.

Additionally, while intraoperative microstimulation studies like this provide valuable insight involving electrophysiological signatures, there are also limitations. We assessed tremor reduction on a less than 10 sec time interval. Any increased or decreased tremor reduction responses to chronic stimulation were unable to be evaluated. Additionally, microstimulation results in a lower population of neuronal activation than macrostimulation.^29^ Macrostimulation is utilized in DBS implantations, but results from these microstimulation studies may allow for the more targeted implantation of long-term macroelectrodes. Although QEI was associated with acute tremor reduction, prospective validation will be necessary to establish it as a mechanistic intraoperative biomarker. It has been established that chronic Vim-DBS can result in tolerance and return of tremor, likely due to long-term plasticity changes within cerebellar networks.^30^ In future works, patients with macrocontacts implanted in sites with high percentages of QEI could be continually evaluated for tremor reduction to assess the effects that long-term plasticity has on clinical outcomes, and whether this neurophysiological signal is protective of habituation.

## Conclusions

A new objective methodology was utilized to classify single-unit extracellular potential recordings as QEI positive or negative. Moreover, the observation of QEI during high frequency stimulation may be used as a mechanistic biomarker of clinical inhibition. These results might pave the way for closed-loop optimization methodologies for tremor disorders.

## References

1. Song P, Zhang Y, Zha M, et al. The global prevalence of essential tremor, with emphasis on age and sex: A meta-analysis. J Glob Health. 11:04028. doi:10.7189/jogh.11.04028

2. Louis ED, Faust PL, Vonsattel JPG, et al. Neuropathological changes in essential tremor: 33 cases compared with 21 controls. Brain. 2007;130(12):3297–3307. doi:10.1093/brain/awm266

3. Paris-Robidas S, Brochu E, Sintes M, et al. Defective dentate nucleus GABA receptors in essential tremor. Brain. 2012;135(1):105–116. doi:10.1093/brain/awr301

4. Strafella A, Ashby P, Munz M, Dostrovsky JO, Lozano AM, Lang AE. Inhibition of voluntary activity by thalamic stimulation in humans: Relevance for the control of tremor. Movement Disorders. 1997;12(5):727–737. doi:10.1002/mds.870120517

5. Milosevic L, Kalia SK, Hodaie M, Lozano AM, Popovic MR, Hutchison WD. Physiological mechanisms of thalamic ventral intermediate nucleus stimulation for tremor suppression. Brain. 2018;141(7):2142–2155. doi:10.1093/brain/awy139

6. Farokhniaee A, McIntyre CC. Theoretical principles of deep brain stimulation induced synaptic suppression. Brain Stimulation. 2019;12(6):1402–1409. doi:10.1016/j.brs.2019.07.005

7. Paraskevopoulos Z, Tian Y, Crompton D, et al. Frequency-dependent inhibition during deep brain stimulation of thalamic ventral intermediate nuclei. J Neurosci. 2026;46(19):e1859252026. doi:10.1523/JNEUROSCI.1859-25.2026

8. Tian Y, Bello E, Crompton D, et al. Uncovering network mechanism underlying thalamic deep brain stimulation. Preprint posted online December 10, 2023. doi:10.1101/2023.12.09.570924

9. Jager P, Moore G, Calpin P, et al. Dual midbrain and forebrain origins of thalamic inhibitory interneurons. eLife. 2021;10:e59272. doi:10.7554/eLife.59272

10. Hooks BM, Mao T, Gutnisky DA, Yamawaki N, Svoboda K, Shepherd GMG. Organization of cortical and thalamic input to pyramidal neurons in mouse motor cortex. J Neurosci. 2013;33(2):748–760. doi:10.1523/JNEUROSCI.4338-12.2013

11. Ambardekar AV, Ilinsky IA, Forestl W, Bowery NG, Kultas-Ilinsky K. Distribution and properties of GABA(B) antagonist [3H]CGP 62349 binding in the rhesus monkey thalamus and basal ganglia and the influence of lesions in the reticular thalamic nucleus. Neuroscience. 1999;93(4):1339–1347. doi:10.1016/s0306-4522(99)00282-1

12. Papavassiliou E, Rau G, Heath S, et al. Thalamic deep brain stimulation for essential tremor: relation of lead location to outcome. Neurosurgery. 2004;54(5):1120–1130. doi:10.1227/01.NEU.0000119329.66931.9E

13. Middlebrooks EH, Okromelidze L, Wong JK, et al. Connectivity correlates to predict essential tremor deep brain stimulation outcome: Evidence for a common treatment pathway. Neuroimage Clin. 2021;32:102846. doi:10.1016/j.nicl.2021.102846

14. Oostenveld R, Fries P, Maris E, Schoffelen JM. Fieldtrip: open source software for advanced analysis of meg, eeg, and invasive electrophysiological data. Computational Intelligence and Neuroscience. 2011;2011:1–9. doi:10.1155/2011/156869

15. Horn A, Kühn AA. Lead-DBS: A toolbox for deep brain stimulation electrode localizations and visualizations. NeuroImage. 2015;107:127–135. doi:10.1016/j.neuroimage.2014.12.002

16. Friston KJ, Ashburner J, Kiebel S, Nichols T, Penny WD, eds. Statistical Parametric Mapping: The Analysis of Funtional Brain Images. First edition. Elsevier/Academic Press; 2007.

17. Avants B, Tustison NJ, Song G. Advanced normalization tools: v1. 0. The Insight Journal. Published online July 29, 2009. doi:10.54294/uvnhin

18. Jenkinson M, Bannister P, Brady M, Smith S. Improved optimization for the robust and accurate linear registration and motion correction of brain images. NeuroImage. 2002;17(2):825–841. doi:10.1006/nimg.2002.1132

19. Woolrich MW, Jbabdi S, Patenaude B, et al. Bayesian analysis of neuroimaging data in FSL. NeuroImage. 2009;45(1):S173–S186. doi:10.1016/j.neuroimage.2008.10.055

20. Ewert S, Plettig P, Li N, et al. Toward defining deep brain stimulation targets in MNI space: A subcortical atlas based on multimodal MRI, histology and structural connectivity. NeuroImage. 2018;170:271–282. doi:10.1016/j.neuroimage.2017.05.015

21. Interactive data visualization tools & software | plotly. Accessed July 26, 2026. https://plotly.com/

22. Chechile RA. A bayesian analysis for the mann-whitney statistic. Communications in Statistics - Theory and Methods. 2020;49(3):670–696. doi:10.1080/03610926.2018.1549247

23. Xu SS, Lee WL, Perera T, et al. Can brain signals and anatomy refine contact choice for deep brain stimulation in Parkinson’s disease? J Neurol Neurosurg Psychiatry. Published online May 19, 2022:jnnp-2021-327708. doi:10.1136/jnnp-2021-327708

24. Ozturk M, Viswanathan A, Sheth SA, Ince NF. Electroceutically induced subthalamic high-frequency oscillations and evoked compound activity may explain the mechanism of therapeutic stimulation in Parkinson’s disease. Commun Biol. 2021;4(1):393. doi:10.1038/s42003-021-01915-7

25. Birdno MJ, Kuncel AM, Dorval AD, Turner DA, Gross RE, Grill WM. Stimulus features underlying reduced tremor suppression with temporally patterned deep brain stimulation. J Neurophysiol. 2012;107(1):364–383. doi:10.1152/jn.00906.2010

26. Holsapple J, Preston J, Strick P. The origin of thalamic inputs to the “hand” representation in the primary motor cortex. J Neurosci. 1991;11(9):2644–2654. doi:10.1523/JNEUROSCI.11-09-02644.1991

27. Essential tremor and the cerebellum. In: Handbook of Clinical Neurology. Vol 155. Elsevier; 2018:245–258. doi:10.1016/B978-0-444-64189-2.00016-0

28. Coenen VA, Sajonz B, Prokop T, et al. The dentato-rubro-thalamic tract as the potential common deep brain stimulation target for tremor of various origin: an observational case series. Acta Neurochir. 2020;162(5):1053–1066. doi:10.1007/s00701-020-04248-2

29. Wu YR, Levy R, Ashby P, Tasker RR, Dostrovsky JO. Does stimulation of the GPi control dyskinesia by activating inhibitory axons? Movement Disorders. 2001;16(2):208–216. doi:10.1002/mds.1046

30. Peters J, Tisch S. Habituation after deep brain stimulation in tremor syndromes: prevalence, risk factors and long-term outcomes. Front Neurol. 2021;12:696950. doi:10.3389/fneur.2021.696950

