## Supplementary for "Inhibitory Evoked Potentials as a Spatially Dependent Intraoperative Marker of Clinical Tremor Reduction"

### Supplementary Tables

| Patient | Sex | Procedure | Data | Direction |
|---|---|---|---|---|
| 1 | Male | Thalamotomy | MER, MRI, accelerometer | Right |
| 2 | Male | Thalamotomy | MER, MRI, accelerometer | Left |
| 3 | Female | DBS | MER, MRI, accelerometer | Left |
| 4 | Male | Thalamotomy | MER, MRI, accelerometer | Left |
| 5 | Male | DBS | MER, accelerometer | Bilateral |
| 6 | Male | DBS | MER, MRI, accelerometer | Left |
| 7 | Male | Thalamotomy | MER, accelerometer | Left |
| 8 | Male | Thalamotomy | MER, accelerometer | Left |
| 9 | Male | DBS | MER, MRI, accelerometer | Left |
| 10 | Male | DBS | MER, MRI, accelerometer | Right |
| 11 | Male | DBS | MER, accelerometer | Right |
| 12 | Male | DBS | MER, accelerometer | Left |

Supplementary table 1: Patient demographics and recording modalities. Each patient's sex, whether they received DBS or a surgical thalamotomy, what data types were recorded, and the side at which the surgical target was on is listed.

### Supplementary methods

#### Human experimental data

Before their involvement in the study, patients provided written informed consent. Additionally, the study protocol was in agreement with the Declaration of Helsinki, conformed to the guidelines set by the Tri-Council Policy on Ethical Conduct for Research Involving Humans, and were approved by the University Health Network Research Ethics Board. Two closely spaced microelectrodes (~600 um apart) were used to both stimulate and record single units. Microstimulation was delivered using variable intensity and symmetric 0.3ms biphasic pulses (150μs cathodal followed by 150 μs anodal). The stimulation frequencies were 100 Hz (n = 26), 200 Hz (n = 88), and 300 Hz (n = 22), and train duration varied between 0.8-5 sec based on the experiment and possible interruption due to side effects. Recordings were sampled at 10 kHz. A total of 136 stimulation trains were analyzed.

#### Detection and classification of quasi-evoked inhibition

To preprocess the data before detection of quasi-evoked inhibition, a third order Savitzky-Golay filter was applied to frame lengths of 0.9 ms to remove noise. Then the three-step artifact removal with quasi-evoked inhibition preservation from Paraskevopoulos et al. 2025 [1] was applied. For each inter-pulse interval, this involved applying an exponential fit to the segment prior to the quasi-evoked inhibition event. Then a linear fit to the duration of the quasi-evoked inhibition event, and finally a second exponential fit to the segment following the quasi-evoked inhibition event. The 0.7 mV threshold was chosen as it was the lowest threshold tested in Paraskevopoulos et al. 2025 [1] that produced similar results. The starting voltage was necessary as this often was the location of a spike and any following positive deflection could be the effect of membrane hyperpolarization. The 40% of pulses classification criteria was chosen because this threshold was chosen as the lowest percentage of quasi-evoked inhibition events following pulses in Paraskevopoulos et al. 2025 [1] was 44.92%.

Even after filtering, a subset of recordings had a much higher level of noise. To prevent recording noise from being misclassified as a quasi-evoked inhibition event, the amplitude threshold was raised to 0.1 mV (the exact threshold used in Paraskevopoulos et al. 2025 [1]) and the percent of pulses with quasi-evoked inhibition threshold was raised to 60% (as the lowest percentage of quasi-evoked inhibition events following pulses in a high noise recording in Paraskevopoulos et al. 2025 [1] was 83.93% and 60% was the median between low and high noise).
